# Enhancer pioneering activity of Wnt/β-catenin signaling

**DOI:** 10.64898/2026.06.12.731851

**Authors:** Tamina Weiss, Anna Nordin, Jente Zijlstra, Pierfrancesco Pagella, Claudio Cantù

**Affiliations:** Department of Biomedical and Clinical Sciences, Division of Molecular Medicine and Virology; Faculty of Medicine and Health Sciences; Linköping University, Sweden; Wallenberg Centre for Molecular Medicine, Linköping University, Sweden; Science for Life Laboratory – SciLifeLab, Linköping University, Linköping, Sweden; Department of Physics, Chemistry and Biology, Division of Chemistry, Faculty of Science and Engineering, Linköping University, Linköping, Sweden

## Abstract

Wnt/β-catenin signaling is widely understood as a molecular switch that activates preprogrammed regulatory states. Whether it also actively reshapes chromatin to establish those states has remained unclear. To address this, we performed time-resolved ATAC-seq and CUT&RUN analyses in HEK293T cells and monitored β-catenin-dependent changes in chromatin accessibility and complex assembly following pathway activation. In addition to conventional Wnt-responsive elements (WREs), which are constitutively accessible and pre-bound by TCF/LEF transcription factors (TFs), we identified a second class of WREs, that are initially inaccessible but become activated upon stimulus. At these latent enhancers, β-catenin functions as a pioneer-like factor together with HDAC1 and CBP, mediating chromatin remodeling and TCF/LEF occupancy. Together, these results extend the current view of Wnt/β-catenin signaling by showing that it can establish regulatory elements *de novo*. Here, β-catenin emerges as the central regulator of transcriptional control and remodeling, providing a framework for the dynamic and context-dependent nature of Wnt-driven gene regulation in development and disease.

## Introduction

Wnt/β-catenin signaling is an evolutionarily conserved signal transduction cascade with pivotal functions throughout embryogenesis, stem cell homeostasis, and regeneration^1^. Central to the signaling cascade is the cytoplasmic regulation of β-catenin by the destruction complex, composed of the tumor suppressors AXIN, APC, CK1α, and GSK3. This complex constitutively phosphorylates β-catenin, thereby priming it for proteasomal degradation and thus maintaining the pathway in an OFF-state. Binding of WNT ligands to the cell-surface FRIZZLED and LRP receptors on signal receiving cells leads to inhibition of the destruction complex. Thus, cytosolic β-catenin accumulates above a critical threshold, enabling its translocation to the nucleus (ON-state)^2^. Here β-catenin coordinates binding of a complex containing transcriptional co-activators including CPB/p300^3,4^, the BAF complex^5–7^, BCL9/9L, and PYGO^8–10^ on so called Wnt-responsive elements (WRE). β-catenin itself lacks intrinsic DNA-binding capacity and therefore relies primarily on interactions with TCF1, TCF7L1, TCF7L2 or LEF1, the members of the TCF/LEF family of transcription factors (TFs)^11^ to bind DNA. TCF/LEF-independent β-catenin-DNA associations have also been reported^12,13^, mediated by for example alternative TFs such as SOX17 during mesoderm patterning^13^.

Despite these insights, our understanding of how incoming Wnt signals are integrated at the chromatin level to generate diverse transcriptional programs remains incomplete^14–17^. One prevailing view is that the cellular context acts as the primary determinant of signaling output: (1) the accessibility of WREs bound by TCF/LEF TFs and (2) presence of co-factors dictate which genes become transcriptionally activated, while β-catenin enables the recruitment of transcriptional co-activators^10,18–20^ (Figure 1A). However, whether β-catenin directly contributes to modulating chromatin states remains unclear. Notably, we have observed instances where β-catenin binds to initially inaccessible chromatin, followed by local chromatin opening^21^. This suggests that not only the preceding chromatin states localize signaling factors^22^, but also the presence of a signaling effector, such as β-catenin, may influence regulatory outcomes.

**Fig. 1.**
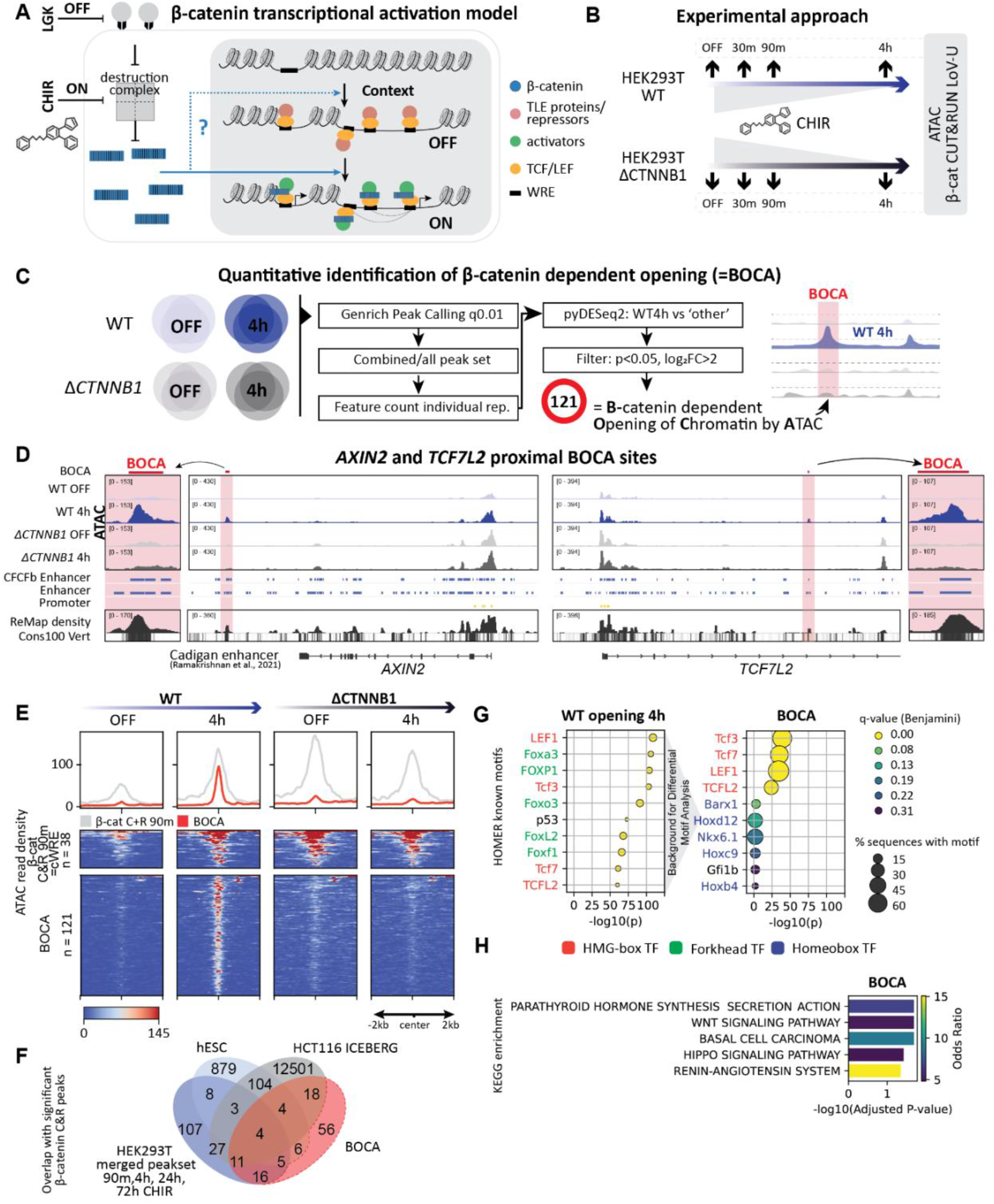
β-catenin dependent chromatin opening. (**A**) Schematic showing nuclear β-catenin as a transcriptional switch that binds TCF/LEF at Wnt-responsive elements, displaces repressors (e.g., TLE), and recruits co-activators to activate transcription. It remains unclear whether β-catenin also directly shapes chromatin accessibility and TCF/LEF positioning. (**B**) Experimental design using time-resolved LGK and CHIR treatments in HEK293T wild-type (WT) and β-catenin knockout (Δ*CTNNB1*) cells, followed by ATAC-seq and CUT&RUN LoV-U profiling. (**C**) Analysis pipeline identifying β-catenin–dependent chromatin opening (BOCA). Peaks were called across replicates (Genrich, q < 0.01, AUC > 200), merged into a unified set, quantified (FeatureCounts), and tested for differential accessibility (pyDESeq2; p < 0.05, log2FC > 2), defining 121 BOCA sites enriched in WT CHIR 4 h. (**D**) ATAC-seq tracks (CPM-normalized, n=3) at representative BOCA loci, including *AXIN2* (Cardigan enhancer) and *TCF7L2* intronic enhancer. Regulatory annotations (enhancers, promoters, CTCF sites), TF binding density(ReMap2022^32^), and vertebrate conservation are shown. (**E**) Average plots (top) and heatmaps (bottom) comparing normalized ATAC-seq signal at β-catenin occupied sites at 90 minutes upon CHIR (see Figure2, cWRE, grey) and BOCA sites (red), ±2 kb around peak centers. For the HEK WT ATAC conditions, previously published data was used^21^. (**F**) Overlap of BOCA sites with β-catenin CUT&RUN peaks from HEK, hESC, and HCT116 datasets^21,29^. (**G**) HOMER motif enrichment identifying transcription factor families associated with WT chromatin opening (left) and BOCA sites(right). Yellow dots indicate significance with q (Bejamini) < 0.05 and −log10(p) > 20 and dot size the percentage of target sequences containing the motif. TF family indicated: HMG-box (red), Forkhead (green), and Homeobox (blue). (**H**) KEGG pathway enrichment of the 200 genes annotated to BOCA sites using GREAT basal plus extension.

These observations prompted us to investigate whether β-catenin may impart functional chromatin state following pathway activation. To this end, we performed time-resolved ATAC-seq (assay for transposase-accessible chromatin using sequencing)^23^ and CUT&RUN-LoV-U (cleavage under targets and release using nuclease-low volume-urea)^24^ assays to quantify the temporal recruitment of the Wnt transcriptional complex components and how this is accompanied by functionally relevant chromatin features. In profiling β-catenin and all TCF/LEF family members in wild-type and *CTNNB1* (human β-catenin gene) knock-out human embryonic kidney 293T cells (HEK WT and HEK *ΔCTNNB1* respectively^12^), we identified two distinct classes of WRE with fundamentally different regulatory properties. We show that β-catenin, despite lacking intrinsic DNA-binding capabilities, can function in a pioneer-like manner to induce enhancer activation. β-catenin employes a TCF/LEF-mediated mechanism of motif-dependent enhancer recognition and combines CBP/HDAC enzymatic activities to progressively remodel the chromatin. These findings indicate that the β-catenin complex functions not only as a signal amplifier, but as an active regulator of chromatin states.

## Results

### Distinct sites require β-catenin for chromatin opening upon Wnt pathway activation

We previously identified genomic regions that gain accessibility upon WNT pathway activation in HEK WT cells (Figure 1A-B)^21^. Cells were first driven into a Wnt-OFF state by treatment with the PORCN inhibitor LGK (10 nM) for 24 hours, followed by exactly 4 hours with GSK3 inhibitor CHIR99021 (10 µM, hereafter CHIR) to induce the Wnt-ON state. GSK3 inhibition has pleiotropic effects, some of which have been shown to be independent from the GSK3-β-catenin axis^25^. Thus, we replicated this experiment in HEK Δ*CTNNB1* cells (Figure 1B; cells from Doumpas et al., (2019)) and generated matching ATAC-seq data at the corresponding time points, enabling direct comparison of Wnt-induced chromatin accessibility in the presence and absence of β-catenin.

To identify sites that are β-catenin dependent during pathway induced chromatin opening (i.e., regions opening in WT but not in Δ*CTNNB1* cells), we peak-called ATAC-seq data from all four conditions using Genrich (p < 0.05, a = 200) leveraging its built-in function to consider all three replicates. Peaks from all conditions were merged into a union peak set which was subsequently used to quantify ATAC-seq signal for each individual replicate and condition using FeatureCount^26^. To find sites that have a significantly higher read-count in the WT condition compared to the rest, we used the Wald-statistics of pyDESeq2 (v.0.5.1) and filtered for significant results (p < 0.05, log2FC > 2). This resulted in a list of 121 sites that we refer to as ‘B -Catenin dependent Opening of chromatin using ATAC’, hereafter BOCA (Figure 1C). Consistently, BOCA sites show opening upon CHIR treatment in WT cells, whereas they remain inaccessible in Δ*CTNNB1* cells (Figure 1D-E). The two representative sites shown in Figure 1D are highly conserved, known enhancers regulating the Wnt output genes *AXIN2*^27^ and *TCF7L2*^28^.

Approximately half of the BOCA sites overlap with previously identified β-catenin binding peaks across multiple biological contexts, including HEK and hESC cells treated with CHIR for 90 minutes, 4h, 12h 24h and 72h^21^, and HCT116 cells^29^. HOMER (v5.1) known motif analysis of regions underlying WT opening and BOCA sites revealed strong enrichment for all four TCF/LEF motifs (Figure 1G). GREAT’s^30^ basal plus extension function annotated the 121 BOCA sites to 200 genes, including *AXIN2, LEF1, TCF7L2* and *DKK1*. KEGG pathway enrichment analysis (Enrichr^31^, whole genome background) resulted in ‘Wnt signaling pathway’ (p-adj = 0.02, OR = 5.05) and ‘Hippo signaling pathway’ (p-adj = 0.03, OR = 4.81) among the top 5 significant hits (Figure 1H). Collectively, these results reveal that there are genomic sites that depend on β-catenin for chromatin opening upon pathway activation, and these sites are well associated to the Wnt transcriptional output, as indicated by underlying motifs and associated genes.

### TCF/LEF require β-catenin to bind the BOCA sites

We tested if, and when, β-catenin becomes associated with the BOCA sites upon pathway activation. We did a replicated time-resolved CUT&RUN series of β-catenin in HEK WT using the Low-Volume Urea protocol (LoV-U)^24^ and indeed found that association to chromatin in BOCA regions increases within the first 90 minutes upon CHIR administration. This association is lost in the knock-out cells (Figure 2A-C, blue tracks).

**Figure 2.**
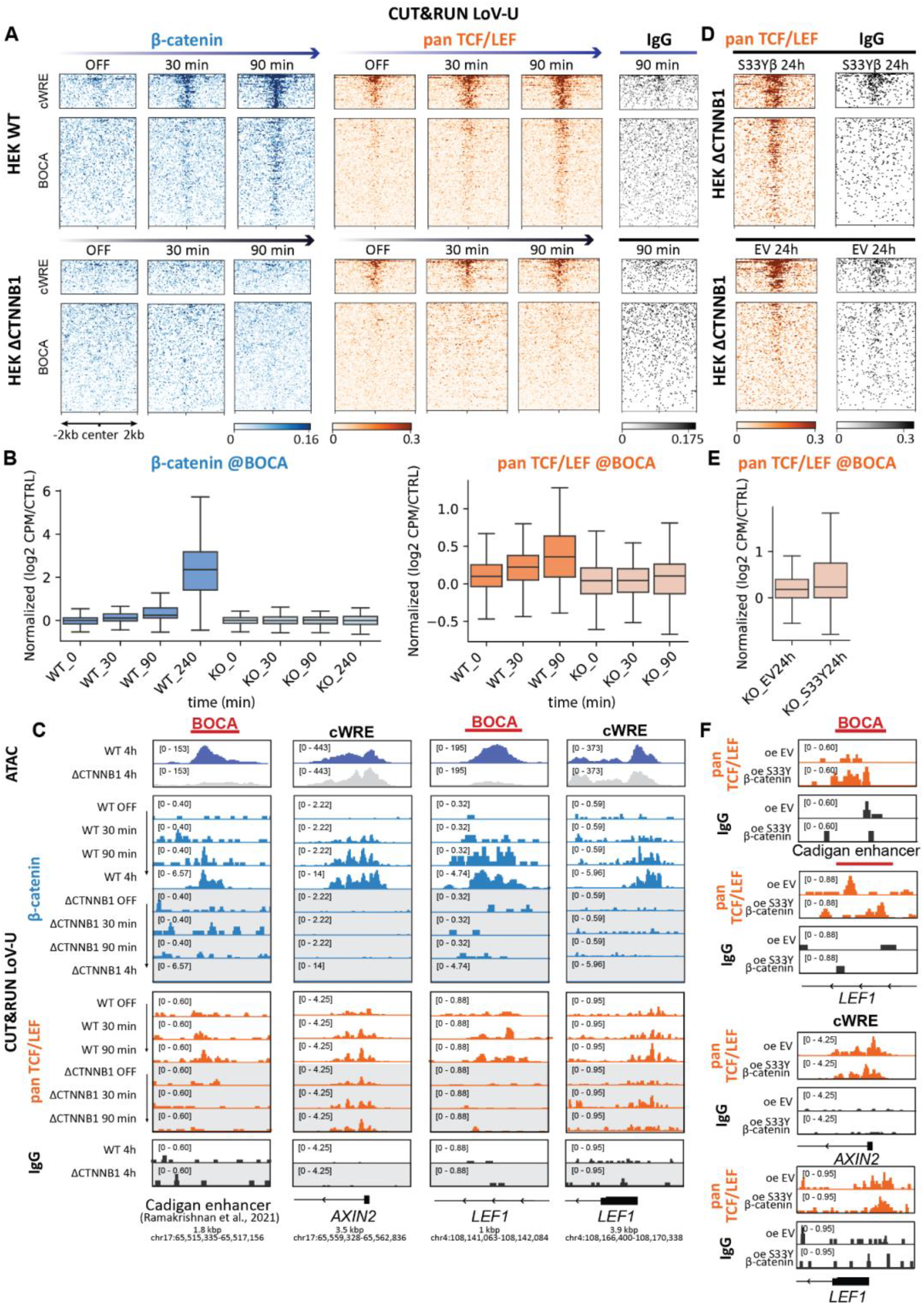
TCF/LEFs need β-catenin for recruitment to BOCA sites. **(A)** Heatmaps comparing CPM-normalized CUT&RUN signal of β-catenin (blue), all 4 members of the TCF/LEF family of TFs (=panTCF/LEF, orange) and an IgG control (black), ±2 kb around WREs (β-catenin occupancy at CHIR 90 min, grey) and BOCA sites (red). Shown are HEK WT and HEK Δ*CTNNB1* at Wnt-OFF (10nM LGK), or upon Wnt activation using CHIR (10µM) for 30, or 90 minutes. (**B)** Quantification of β-catenin (blue) or panTCF/LEF (orange) CUT&RUN features at each BOCA site and sample, as log2[(sample CPM/IgG CPM) +1]. Boxes indicate IQR, median shown as a horizontal line; whiskers extend to 1.5x IQR. KO denotes HEK Δ*CTNNB1* (lighter shades), time in minutes upon CHIR stimulation. **(C)** Signal tracks of CPM normalized, merged ATAC-seq (upper squares) or β-catenin (blue), panTCF/LEF (orange) and IgG (black) CUT&RUN samples in HEK WT and HEK Δ*CTNNB1* (grey boxes) at several time points upon WNT activation at two representative BOCA sites (red bars, *AXIN2* cardigan enhancer and *LEF1* enhancer) and conventional WRE (*AXIN2* promoter and *LEF1* promoter). (**D)** Heatmaps (bottom) of CPM normalized panTCF/LEF (orange) and IgG control (black) CUT&RUN signal derived from HEK Δ*CTNNB1* cultured in Wnt OFF (LGK10nM) that overexpress either an empty vector control (oe EV) or a constitutively active β-catenin harboring a S33Y mutation (oe S33Y β-catenin) for 24h. (**E)** panTCF/LEF (orange) CUT&RUN features at BOCA sites, quantified for the samples from D. (**F)** Exemplary signal tracks from C shown at representative genomic loci.

Since we observed that BOCA sites are enriched for TCF/LEF motifs, we were curious whether TCF/LEF TFs can associate to these motifs even within closed chromatin and thereby initiate changes in chromatin accessibility. Such binding, as the current tenet in the field would imply, should occur independently of β-catenin. We first focused on high-confidence β-catenin binding sites found at 90 minutes (MACS2 on two replicates at q < 0.01 over the IgG control (n = 38)). In a similar time-resolved CUT&RUN series targeting all TCF/LEF TFs, we found that, as expected, TCF/LEF factors bind to these WREs (Figure 2A, orange upper boxes). These WREs include well known Wnt response loci such as the *AXIN2* promoter (Figure 2C), and TCF/LEFs associate to them irrespective of pathway activation (in both Wnt ON and OFF) and independently from β-catenin (both in HEK WT and Δ*CTNNB1*). As this behavior matches the expectation from current knowledge, we refer to these genomic regions as Conventional WREs (C-WREs). To our surprise, and in stark contrast, at BOCA sites TCF/LEF occupancy was not detected in the Wnt OFF state and increased within the first 90 minutes of pathway activation (Figure 2B). Of note, TCF/LEF association at BOCA sites, but not at C-WREs, was lost in the absence of β-catenin (orange Figure 2A-B, tracks with gray background in Figure 2C). Testing all TCF/LEFs together (pan-TCF/LEF) or each member individually confirmed this observation (Supplementary Figure 2B-C).

The inability of TCF/LEF to occupy BOCA sites in the absence of β-catenin was restored upon overexpression of a constitutively active β-catenin mutant (S33Y), which evades proteasomal degradation (Figure 2D, 2E and 2F). This rescue underscores that β-catenin is not only necessary, but also sufficient in enabling TCF/LEF engagement at these sites.

### BOCA sites are (poised) Wnt-dependent enhancers

Genomic annotation using HOMER^33^ further revealed that none of the BOCA sites are a promoter; they are instead predominantly located in intergenic and intronic regions (Figure 3B). Their conservation and transcription factor binding, as indicated by enrichment in PhastCons100 vertebrates and ReMAP2022^32^ signal respectively (Supplementary Figure 3), suggest that these regions are cis-regulatory elements, or enhancers, across systems. Enhancers are characterized by specific histone modifications that reflect their regulatory state. Mono-methylation of histone H3 lysine 4 (H3K4me1) is broadly associated with both poised and active enhancers, whereas presence of acetylation of histone H3 lysine 27 (H3K27ac) distinguishes active enhancers from poised or inactive ones, which are typically marked by H3K4me1 alone^34– 36^. Because we found that β-catenin and TCF/LEF association to previously closed chromatin is followed by chromatin opening, we sought to determine whether these sites acquire corresponding changes in the local epigenetic landscape.

**Figure 3.**
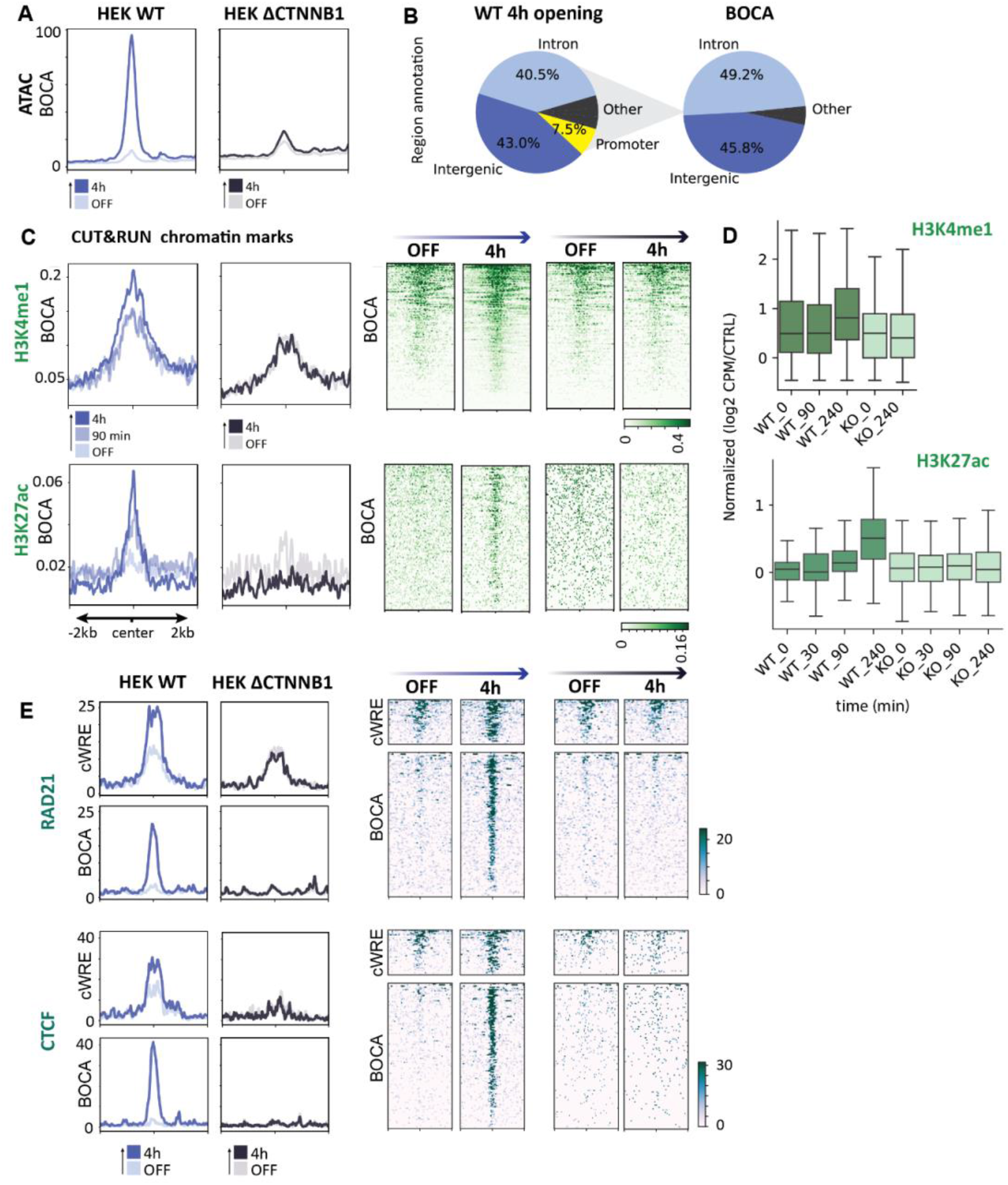
β-catenin dependent activation of enhancers. **(A)** Average signal at BOCA sites plots comparing CPM normalized ATAC-seq samples comparing HEK WT (purples) and HEK Δ*CTNNB1 (greys)* at Wnt-OFF, or upon Wnt activation using CHIR for 4 hours (n=3). (**B**) Genomic annotation of ATAC peaks from all WT opening events upon CHIR 4 hours (Supplementary Figure 1) and BOCA sites, categorized as promoter, intronic, intergenic, or other regions using HOMER^33^. (**C)** Average plots (left) and heatmaps (right) showing CPM normalized presence of the activating chromatin marks H3K4me1 and H3K27ac (n=2) across all BOCA sites assayed by CUT&RUN. Signal is derived from HEK WT (purples) and HEK Δ*CTNNB1* (greys) at WNT-OFF, Chir 90 minutes or CHIR 4h (shown with increasing color intensities). (**D)** H3K4me1 (top) and H3K27ac (bottom) CUT&RUN features at BOCA sites, counted per sample and normalized as log2[(sample CPM/IgG CPM) +1]. Boxes indicate the interquartile range with the median shown as a horizontal line; whiskers extend to 1.5x IQR. KO denotes HEK Δ*CTNNB1* (lighter shades), time in minutes upon CHIR stimulation. (**E)** Average plots (left) and heatmaps (right) showing replicated, normalized chromatin association of RAD21 (member of the cohesion complex) and CTCF, across standard WRE and BOCA sites across cell lines and times upon pathway activation (n=3, data from^37^).

At BOCA sites in HEK-WT cells, 4 hours of CHIR treatment led to increased signal for H3K4me1 and H3K27ac. This pathway-dependent increase was not observed in the Δ*CTNNB1* cells (Figure 3C-D). Of note, the average H3K4me1 CUT&RUN profile in untreated HEK WT cells was already above background, indicating that enhancers are poised but inactive in closed chromatin prior to pathway activation35. Nevertheless, in the absence of β-catenin, this poised enhancer state is insufficient for priming the chromatin both for activation and opening, as evidenced by their lack of CHIR-induced gain in H3K4me1 and H3K27ac CUT&RUN signal (Figure 3C-D), and ATAC signal (Figure 3A), respectively.

Active, open enhancers enable binding of TFs and the association of architectural proteins, including cohesins and CTCF, that foster enhancer–promoter looping and the stabilization of the transcriptional machinery. Notably, BOCA sites are only occupied by CTCF and RAD21 (a cohesin subunit) upon pathway activation, demonstrating their functional importance in shaping gene regulatory networks (Figure 3E). In Δ*CTNNB1* cells, loss of enhancer activity and chromatin accessibility coincides with disrupted interactions, suggesting profound consequences for 3D genome organization. Accordingly, our group has recently demonstrated that CTCF binding repositions upon Wnt pathway activation, and that this redistribution is largely dependent on β-catenin, with clear structural consequences for 3D genome organization as measured by CTCF HiChIP37.

These data show BOCA sites display the features of poised/latent enhancers that require β-catenin for activation. At BOCA sites, in HEK-WT cells, Wnt signaling triggers H3K4me1 and H3K27ac accumulation in parallel with chromatin opening, recruitment of transcription factors and the architectural proteins CTCF and RAD21, which likely influences the expression of Wnt target genes (Supplementary Figure 2D). In Δ*CTNNB1* cells, these events fail to occur, leaving chromatin closed and interactions disrupted, highlighting the essential role of β-catenin in converting poised BOCA enhancers into active regulatory elements.

### Chromatin remodeling dynamics distinguish BOCA sites from C-WREs

Histone acetylation of nucleosomes plays a central role in controlling chromatin accessibility. Early studies identified among the β-catenin co-factors the histone acetyltransferases CBP/p300^38^ and the histone deacetylase HDAC1^39^. We tested whether changes in the epigenetic landscape at BOCA sites following pathway activation might be due to active recruitment of these enzymes via β-catenin. CUT&RUN experiments targeting either CBP/p300 or HDAC1 revealed a modest, time-dependent increase in their signal within the first 90 minutes of treatment. Interestingly, also CBP and HDAC1 recruitment appeared to be β-catenin dependent as they failed to accumulate in Δ*CTNNB1* cells (Figure 4A-B, 4D-E lower panels).

**Figure 4.**
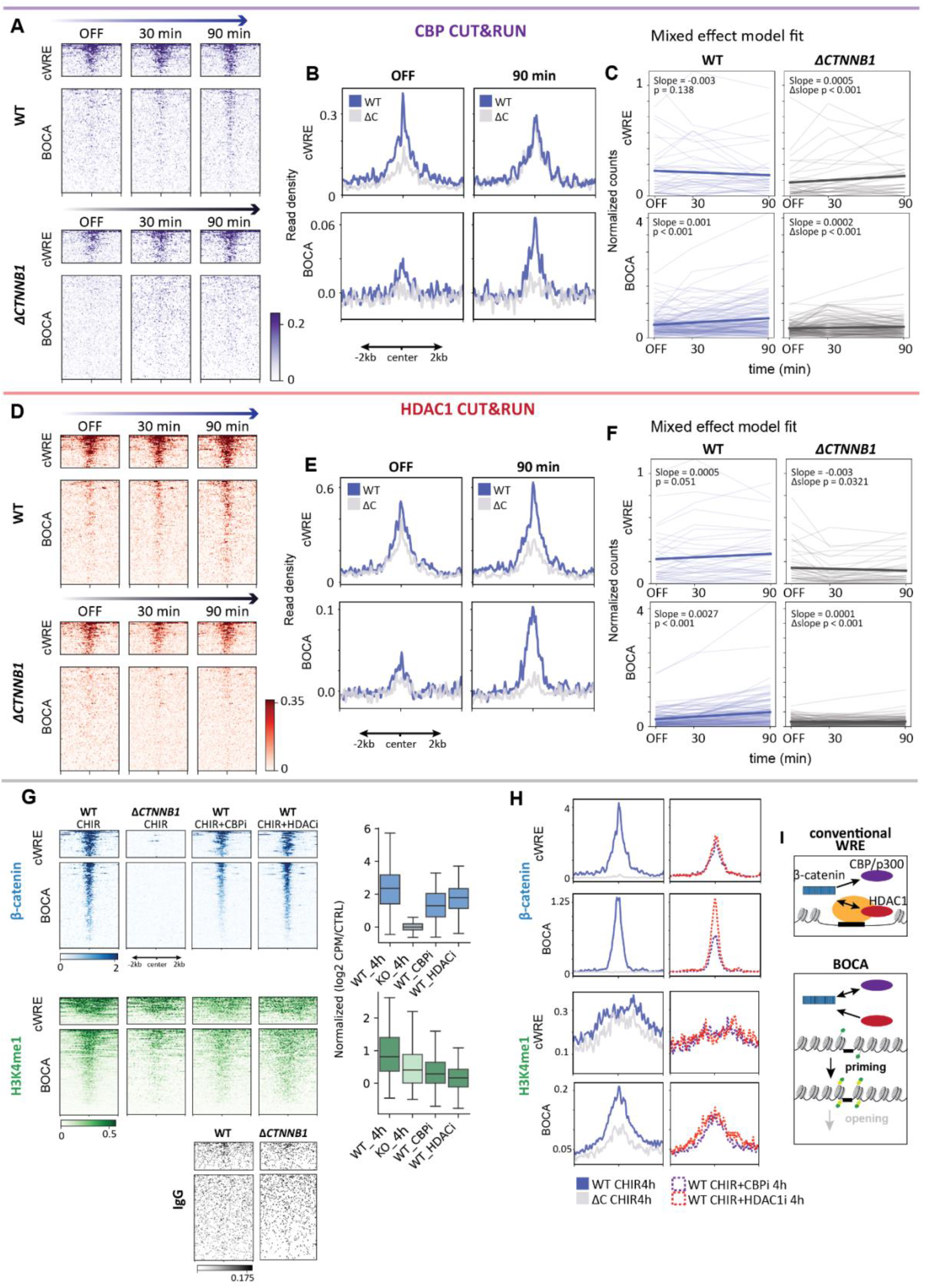
β-catenin associates with CBP/p300 and HDAC1 at BOCA. (A–B) Heatmap (A, left) and average profile (B, right) of CPM-normalized CBP CUT&RUN signal across ±2 kb regions cantered on ‘conventional WREs’ (upper panels) and BOCA sites (lower panels) during the first 90 min of CHIR treatment. Signals from HEK WT (purple) and HEK *ΔCTNNB1* (grey) are shown over time. **(C**) Linear mixed-effects modeling of CBP signal dynamics. Bold lines represent global model fits for HEK WT (purple) and HEK *ΔCTNNB1* (grey) across time, based on CPM-normalized counts at conventional WREs and BOCA sites, with peak identity included as a random effect. Shaded lines indicate individual site trajectories. Annotated slopes reflect time-dependent changes; p-values correspond to the effects of time and the time × genotype interaction. Model: *signal ∼time + genotype + time:genotype + (1*|*peak)*. **(D–E)** As in A–B, but showing CPM-normalized HDAC1 CUT&RUN signal. **(F)** As in C, but modeling HDAC1 signal dynamics across time. **(G)** Left: Heatmaps of CPM-normalized CUT&RUN signal for β-catenin (blue), H3K4me1 (green), and IgG (black) following CBP/p300 or HDAC1 inhibition, across ±2 kb regions centered on conventional WRE (upper panels) and BOCA sites (lower panels). Right: Quantification of CUT&RUN signal at individual BOCA sites, expressed as log2[(CPM / control CPM) + 1] per sample. Boxes represent the IQR with median; whiskers extend to 1.5× IQR. CBP inhibition (CBPi): 10 µM CHIR + 5 µM A-485 for 4 hours; HDAC1 inhibition (HDAC1i): 10 µM CHIR + 1 mM sodium butyrate for 4 hours. (**H)** Average signal profiles corresponding to panel

Linear mixed-effects modelling revealed distinct dynamics of CBP/p300 and HDAC1 occupancy at BOCA sites compared to C-WREs (Figure 4C, 4F). Signal (CPM normalized counts at sites) was modeled using time, genotype, and their interaction as fixed effects, and sites as a random intercept (signal ∼time + genotype + time:genotype + (1∣site)) (Supplementary Figure 4). At BOCA sites, both CBP and HDAC1 showed significant increases in signal over time in HEK WT (CBP: β = 0.001, p < 0.001; HDAC1: β = 0.003, p < 0.001). Significant negative time-by-KO interactions for both factors (CBP: β = −0.001, p < 0.001; HDAC1: β = −0.003, p < 0.001) indicate that their time-dependent recruitment is attenuated in the absence of β-catenin, with a stronger effect observed for HDAC1. In contrast, at C-WREs in HEK WT, the time effect was not significant for CBP (p = 0.138) and for HDAC1 it was lower (β = 0.001, p = 0.051). This model suggests that BOCA sites are characterized by more pronounced and β-catenin-dependent temporal recruitment of both CBP and HDAC1, whereas C-WREs, in line with previous model of Wnt signaling^10^, appeared to be constitutively occupied with limited dynamics.

As shown above, both CBP/p300 and HDAC1 required β-catenin to associate with BOCA enhancers and initiate chromatin remodeling and activation. We next asked whether β-catenin itself depends on these chromatin remodelers to associate with these sites. To address this, we used the small-molecule inhibitors A-485 (5 µM) and sodium butyrate (1 mM), that specifically inhibit CBP/p300^40^ and HDACs^41^, respectively. Concomitant treatment with these inhibitors during 4 hours of Wnt pathway activation (CHIR 10 µM) revealed site-specific mechanistic effects (Figure 4G-H). At C-WREs, inhibition of either CBP/p300 or HDAC1 reduced β-catenin association compared to CHIR-only treated controls. In contrast, at BOCA sites, β-catenin association was reduced only upon CBP/p300 inhibition, while HDAC1 inhibition had no effect. Interestingly, enhancer priming, as measured by H3K4me1 levels, was disrupted by inhibition of either CBP/p300 or HDAC1, suggesting that both remodelers contribute to establishing the active chromatin state even if only CBP/p300 is strictly required for β-catenin recruitment at BOCA sites.

Taken together, these results indicate a site-specific interplay between β-catenin and chromatin remodelers. At BOCA sites, β-catenin drives the dynamic recruitment of CBP/p300 and HDAC1, with CBP/p300 also being required for β-catenin association. In contrast, C-WREs show constitutive occupancy with limited temporal dynamics, although both remodelers still support β-catenin binding. This suggests that C-WREs exist in a more pre-established chromatin state, while BOCA sites are Wnt-dependent enhancers pioneered by the transcriptional complex tethered to the DNA by β-catenin.

## DISCUSSION

The standard model of Wnt signaling-dependent transcription posits that, in the absence of WNT ligands, TCF/LEF transcription factors occupy WREs and recruit corepressors such as Groucho/TLE, maintaining a transcriptionally repressed yet accessible chromatin state, in part through HDAC1 activity^18–20^. Upon pathway activation, β-catenin is recruited to these pre-bound sites, promoting assembly of a higher-order enhanceosome that drives activation and chromatin modifications such as increased H3K27 acetylation^10,42^. Within this framework, β-catenin is thought to function primarily as an activator at already accessible sites, reinforcing pre-existing regulatory configurations. In this work, we challenged this view.

We set out to map the genome-wide regulation of the Wnt/β-catenin dependent transcription over a stimulation time course. The data we obtained uncovers two different modes of action of this pathway. The first aligns with the standard model, according to which accessible WREs are constitutively bound by the TCF/LEFs, and only after Wnt pathway activation are they engaged by β-catenin and its cofactors to foster chromatin remodeling and transcriptional activation. The second—and unexpected—involves a subset of latent regulatory regions, that we call BOCA. These are a mechanistically distinct class of WREs that extend the prevailing model; they are inaccessible and lack detectable TCF/LEF binding prior to stimulation, yet acquire both features upon Wnt activation, in a β-catenin dependent manner. Importantly, our data demonstrates that Wnt signaling, via β-catenin, can induce *de novo* enhancer activation. Similar activation of latent enhancers has been observed in differentiated cells of the immune system responding to inflammatory cues, suggesting that inducible enhancer formation may represent a general mechanism of stimulus-responsive gene regulation^43,44^.

Recent work provides a potential basis for the process we have identified, by showing that β-catenin functions as molecular adapter for the BAF remodeling complex^7^ and that pathway activation promotes TRIP12-dependent ubiquitylation of BRG1 (SMARCA4), enhancing its interaction with β-catenin and facilitating recruitment of the SWI/SNF complex to Wnt target loci^45^. While this mechanism explains how chromatin remodeling may be executed, it does not address how specific loci are selected. Notably, BOCA sites are not pre-bound by TCF/LEF factors. Instead, TCF/LEF recruitment occurs concomitantly with chromatin opening and depends on β-catenin. These sites are also strongly enriched for TCF/LEF motifs, while lacking signatures of other established pioneer factors, in contrast to cWRE that display FOX motifs. Together, these observations place β-catenin functionally upstream of TCF/LEF at BOCA sites and support a role for β-catenin in enabling chromatin engagement. Our data supports a model in which β-catenin, by interacting with the chromatin associating enzymes CBP and HDAC1, can remodel chromatin and render it accessible for TCF/LEF-mediated scanning and motif recognition. This interpretation would entail that not only proteins, but complexes may act as chromatin pioneers. Specifically, the β-catenin-dependent complex exploits the TCF/LEF partners to provide DNA-binding capacity and CBP/HDAC1 moieties to remodel the chromatin and explore the underlying transcription factor binding signatures. Under a functional definition, where pioneering activity refers to the establishment of chromatin competence for transcription factor binding 35, our findings support such a role for the Wnt-dependent transcriptional complex. β-catenin may operate analogously to non-DNA-binding cofactors described in nuclear receptor signaling (GPS2 and NCoR/SMRT regulate accessibility for the signal-dependent transcription factors LXR, PPARγ, and GR^46–50^), or to the non-DNA-binding domain of pioneer proteins like SOX2^51^. These systems demonstrate that cofactors/domains can actively establish permissive chromatin states without directly binding DNA, and that interacting proteins act as individual parts of a functional chromatin-pioneering unit.

Beyond local chromatin remodeling, our findings also implicate higher-order genome organization in Wnt-mediated gene regulation. Previous studies have suggested that enhancer-promoter interactions at Wnt target genes may be pre-established, facilitating rapid transcriptional responses upon pathway activation^52^, but additional chromatin looping has been observed in Wnt-driven positive feedback circuits under high CHIR conditions^52^. Consistent with this, Wnt signaling has been shown to reshape three-dimensional chromatin organization ^37,53^, in part through dynamic rearrangement of CTCF binding^37^. Disruption of CTCF binding at enhancers associated with DKK1 and AXIN2 significantly alters target gene expression, and notably, both enhancers correspond to BOCA sites. In line with these observations, we find that recruitment of CTCF and RAD21 to BOCA enhancers is β-catenin–dependent.

Together, our findings refine and complete the prevailing view of Wnt/β-catenin signaling by showing that its transcriptional output extends beyond activation of pre-existing enhancers to include *de novo* formation of regulatory elements within previously inaccessible chromatin. β-catenin acts not only as a co-activator but as a central orchestrator of chromatin remodeling, transcription factor recruitment, and genome organization. This dual mode of action provides a framework for how Wnt signaling achieves both robustness and flexibility in gene regulation, with broader implications for signal-dependent transcription in development and disease.

## Acknowledgments

We thank all the members of the Cantù lab for the continuous scientific input. The computations and data handling were enabled by resources provided by the National Supercomputer Centre (NSC), funded by Linköping University. We acknowledge the Core Facility at the Faculty of Medicine and Health Sciences, Linköping University for providing sequencing infrastructure and assistance.

## Funding

The Cantù lab is supported by Grants from the Swedish Research Council, Vetenskapsrådet (2021–03075, 2023-01898 and 2025-02369), Cancerfonden (21 1572 Pj and 24 3487 Pj), Additional Ventures (USA) (SVRF2021-1048003), Linköping University and LiU/RÖ Cancer, the Knut och Alice Wallenbergs Stiftelse and SciLifeLab. C.C. is Fellows of the Wallenberg Centre for Molecular Medicine (WCMM) and Group Leader at SciLifeLab and receives generous financial support from the Knut and Alice Wallenberg Foundation.

### Author contributions

T.W., C.C. and P.P. conceived the study. T.W. performed the experiments, curated and analyzed the data, and prepared the figures. A.N., and J.Z. performed preliminary experiments and gave critical scientific and methodological input. C.C. funded and administered the project. C.C. and P.P. supervised the project. T.W. wrote the original draft. C.C edited the original draft. All authors reviewed the manuscript.

## Availability of Data

Raw and processed datasets generated during this study have been deposited ArrayExpress with the accession numbers E-MTAB-16975 (ATAC-seq) and E-MTAB-17186 (CUT&RUN).

## Methods

### Experimental model and cell culture

HEK293T cells WT and ΔCTNNB1 were cultured in DMEM (41966-029, Gibco – Thermo Fisher Scientific) supplemented with 10% Fetal Bovine Serum (12133C, Sigma-Aldrich) and 10 U/ml Penicillin-Streptomycin (15276355, Gibco - Thermo Fisher Scientific). Cells were passaged by incubation in Trypsin EDTA 0.25% (25-200-056, Thermo Fisher Scientific). For all experiments cell passage 13-16 were used.

### Cell treatments

Time-resolved Wnt activation experiments using LGK (S7143, Selleck Chemicals) and CHIR99021 (SML1046, Sigma-Aldrich) were performed as described in Pagella et al. 21. Briefly, 48 hours post-passage, media was supplemented with 10 nM LGK. After 24 hours, cells were treated with 10 µM CHIR99021 for 4 hours. 90, 60, or 30 minutes, or maintained in 10 nM LGK (Wnt-OFF). All conditions were detached simultaneously.

For CBP/p300 or HDAC inhibition, cells were pre-cultured with LGK as described above. After 24 hours, cells were switched to media containing 10 µM CHIR99021 (Wnt-ON) or 10 nM LGK (Wnt-OFF), supplemented with either 5 µM A-485 (CBP/p300 inhibitor54) or 1 mM sodium butyrate (broad HDAC inhibitor41) for 4 hours.

### ATAC sequencing

For HEK293T WT CHIR 4 hours, data was reused from our previous publication (^21^, ArrayExpress E-MTAB-12076). LGK or CHIR 4 hours HEK293T Δ*CTNNB1* data was derived from cells cultured exactly as described in the publication. n = 3 samples, with 5×10^4^ cells per sample were processed for ATAC-seq according to previously published protocols^23^. Briefly, cells were washed with cold PBS, nuclear extracted in ice cold lysis buffer (Tris-HCl pH 7.4 [10 mM], NaCl [10 mM], MgCl_2_ [3 mM], IGEPAL CA-630 [0.1%]), and pelleted. Pelleted nuclei were incubated in 50 μL transposition reaction mix (20034210, Illumina) at 37°C for 30 minutes and the tagmented DNA was purified using QUAGEN Minielute PCR purification. For Library preparation, transposed DNA fragments were amplified for 13 cycles in the presence of Custom Nextera PCR primers^55^ using the NEBNext High-Fidelity 2x PCR Master Mix (Cat. #M0541, New England Biolabs). Libraries were purified using the High Pure PCR Production Purification Kit (11732676001, Roche/ Sigma-Aldrich). Libraries were validated on an Agilent 2100 and quantified using qPCR. Libraries were sequenced on the Illumina NovaSeq 6000 S4 flowcell with PE150 according to results from library quality control and expected data volume.

### ATAC data analysis

Sequencing quality was assessed using FastQC (v0.11.5) with default settings. Adapter trimming and quality filtering were performed using *bbduk* from the BBTools suite (v39.01) in paired-end mode. Adapter removal used the parameters *ktrim=r k=23 mink=11 hdist=1 tpe tbo*, and low-quality bases were trimmed using *qtrim=l trimq=30*, removing reads with Phred scores < 30. Reads were aligned to the GRCh38 reference genome using Bowtie2 (v2.4.5 ^56^) in paired-end mode with the *--very-sensitive-local* setting. Resulting SAM files were converted to BAM format, sorted by genomic coordinates, and indexed using Samtools (v1.22 ^57^). Bam files were used as input for the Genrich (v.0.6.1) peak calling algorithm, levering its built-in function to consider n = 3 replicates. Bedtool’s (v.2.30.0) *intersect* was used to identify HEK WT opening events, defined as the unique HEK WT CHIR 4 hours peaks, not overlapping with the HEK WT LGK peaks. Known motifs enriched in the resulting peak sets were analyzed using HOMER’s (v.4.11) *findMotifsGenome*.*pl*, putting the whole genome as background. For visualization, replicates were merged using SAMTools (v1.22) *merge* and genome-wide coverage tracks as BigWig format were generated using DeepTools’s (v.3.5.6) *bamCoverage* with CPM normalization and *-bin*10 *--extendReads --centerReads --ignoreDublicates*.

### BOCA identification pipeline

Peaks concordant across three replicates were called with Genrich (q < 0.05, AUC > 200, v.0.6.1) for all conditions (HEK293T_WT_LGK, HEK293T_WT_CHIR4h, HEK293T_ Δ*CTNNB1*_LGK and HEK293T_ Δ*CTNNB1*_CHIR4h) and merged into a unified peak set. Fragment counts for all peaks within this union peak set were quantified for each replicate and condition using FeatureCounts^26^. Differential accessibility between HEK293T_WT_CHIR4h and all other conditions was statistically assessed with pyDESeq2 (v.0.5.1) using Wald test statistics and results were filtered for p < 0.05 and log_2_FC > 2. Known motif enrichment in the resulting peak sets was analyzed using HOMER’s (v.4.11) *findMotifsGenome*.*pl*, with the HEK WT CHIR4h dataset as background. Peaks were annotated to genomic features using *annotatePeaks*.*pl*. Gene associations were assigned using GREAT (v4.0.4^30^) with the basal plus extension model (5 kb upstream and 1 kb downstream for proximal regions, and up to 1000 kb for distal regions, including curated regulatory domains), based on hg38 as whole genome background. Gene lists were associated to KEGG biological pathways using Enrichr^31^.

### CUT&RUN LoV-U

Cells were cultured as described above, and all samples were processed using a modified version of the CUT&RUN LoV-U protocol ^24^. Briefly, cells were harvested by trypsinization, washed with PBS, and subjected to three washes with nuclear extraction buffer (HEPES-KOH pH-8.2 [20 mM], KCl [10 mM], Spermidine [0.5 mM], IGEPAL [0.05%], Glycerol [20%], 1xPMSF). For each sample, 250,000 nuclei were bound to 10 µL magnetic ConA beads (Cell Signaling Technology #93569) for 15 minutes at 4 °C with rotation, using the manufacturer’s activation buffer and instructions. Nuclei bound beads were washed with wash buffer (Hepes pH 7.5 [20 mM], NaCl [150 mM], Spermidine [0.5 mM], 1xPMSF) and incubated 5 minutes with wash buffer containing EDTA [0.2 mM], before they were incubated over-night in wash body containing EDTA [0.2 mM], BSA [0.01%], Digitonin [0.025%] and 1:100 antibody. For the panTCF condition, antibodies against LEF1, TCF1, TCF7L1 and TCF7L2 were mixed at equal ratios of 0.8 µL per antibody. For a list of all used antibodies, please refer to table 1. The next day, nuclei were washed five times and incubated 45 minutes at 4 °C in wash buffer containing digitonin [0.025%] and pAG-MNase [0.6 μg/ml]. After additional five washes, nuclei were incubated in 50 µL wash buffer containing CaCl2 [2 mM] for exactly 30 minutes at 4 °C for digestion. Digestion was terminated by adding 3 µL of a 250 mM EDTA/EGTA mix, and the initial fragment release was initiated by the addition of 2 µL NaCl [5 M] per 50 µL sample, followed by incubation at 37 °C for 30 min. This initial release was set aside, and nuclei were incubated in 50 µL urea stop buffer (NaCl [100 mM], EDTA [2 mM], EGTA [2 mM], IGEPAL [0.5%], Urea [8.8 M]) and incubated 30 minutes at RT for a secondary release. Both releases were combined and fragments were purified using phenol chloroform.

**Table 1.** Antibodies and suppliers used for CUT&RUN.

| Target | Manufacturer | Company |
| --- | --- | --- |
| β-catenin | ABIN2855042 | Antibodies Online |
| LEF1 | ABIN1680678 | Antibodies Online |
| TCF7 (TCF1) | ABIN5620945 | Antibodies Online |
| TCF7L1 (TCF3) | ABIN6972849 | Antibodies Online |
| TCF7L2 (TCF4) | CS C48H11 | Cell Signaling Technology |
| CBP | ABIN2854987 | Antibodies Online |
| HDAC1 | ABIN2854776 | Antibodies Online |
| H3K4me1 | CST-5326S | Cell Signaling Technology |
| H3K27ac | ABIN6971893 | Antibodies Online |

### Library preparation and sequencing

Library preparation was performed using the KAPA EvoPrep Kit (Roche, #10096039001) according to the manufacturer’s instructions. For end repair and A-tailing, 14 µL of DNA was used in 0.4x reaction volumes. Adapter ligation was carried out in 0.4x reaction volumes using 0.15 µM KAPA Dual-Indexed adapters, followed by a post-ligation cleanup with Mag-Bind TotalPure NGS beads at a 1x ratio. Libraries were resuspended in 10 mM Tris-HCl (pH 8.0) and amplified using 0.5x reaction volumes. A post-amplification cleanup was performed with 1x beads. Libraries were size-selected by electrophoresis on a 2% E-Gel EX agarose gel (Invitrogen, #G402022) for 10 minutes using the E-Gel Power Snap Electrophoresis System. DNA fragments between 150 and 500 bp were excised and purified using the GeneJET Gel Extraction Kit (Thermo Scientific, #K0691). Libraries were quantified using the Qubit High Sensitivity DNA kit (Thermo Scientific, #Q32854), pooled, and sequenced as 50 bp paired-end reads on a NextSeq 2000 (Illumina) using the NextSeq 1000/2000 P2 XLEAP-SBS Reagent Kit (100 cycles; Illumina, #20100987).

### CUT&RUN data analysis

Paired-end sequencing reads were processed using a standard pipeline. Raw FASTQ files were first subjected to adapter trimming using *BBDuk* (BBTools suite v39.01). Both read pairs were filtered against built in adapters and artifact reference sequences, as well as custom low complexity sequences (poly-AT, poly-G and ployC-stretches). Trimmed reads were then aligned to the human genome (GRCh38/hg38) using Bowtie2 (v.2.4.5) in local alignment mode with the *very-sensitive-local* preset. Only properly paired reads were retained by disabling mixed and discordant alignments, and dovetail alignments were allowed. The insert size range was restricted to 0-500bp. Unaligned reads were excluded from downstream processing. Resulting SAM files were converted to BAM files using SAMtools (v.1.11), followed by mate fixing. PCR duplicates were identified and removed using the *markup* function of SAMtool. To improve read quality, reads mapping to mitochondrial DNA and genomic blacklisted regions were removed using BEDTools (v.2.30.0*) intersect*. The filtered BAM files were subsequently sorted and indexed using SAMTools.

Feature-level read counting was then performed independently for each individual sample, using featureCounts^26^ and a SAF transformed version of the BOCA peak annotations. For all individual BAM files, paired-end reads were counted with stringent filtering options, including fragment-based counting and exclusion of ambiguous or low-quality mappings, ensuring consistent quantification across conditions. To enable normalization, library sizes were calculated for each BAM file by counting properly mapped reads using SAMtool *view -c -F 260*. These counts were paired with sample identifiers to generate per-sample sequencing depth tables for both experimental and control groups. In Python (JupyterLab) using *pandas*, a normalization step was performed, where both matching control (IgG) and sample count matrices were loaded along with their corresponding library sizes. For each genomic feature (e.g., peak), raw counts were converted into CPM using the respective library sizes. A log2-transformed ratio was then computed between sample and control CPM values using a pseudo-count of 1 to avoid zero inflation, producing a normalized enrichment score for each feature across all samples. After checking for consistency, replicates were averaged per biological sample.

For visualization, n = 2 replicates were merged using SAMTools *merge* and genome-wide coverage tracks as BigWig format were generated using deepTools (*bamCoverage*) using CPM normalization and *-bin*10 *--extendReads --centerReads --ignoreDublicates*. Signal tracks were visualized using IGV. Heatmaps and average plots of peak lists of interest (in .bed format) were generated first using DeepTools (v.3.5.6) *computeMatrix* with *--referencePoint center* and *--sortRegions descend*. Matrices were subsequently plotted using the functions *plotHeatmaps* and *plotProfile*, respectively. To define standard WRE in this context, we used the built-in function of MACS2 (v.2.2.6) to peak call individual β-catenin 90 minutes CHIR replicates over the IgG control with *-f BAMPE --keep-dup all –SPMR* as parameters. Peaks from replicates were intersected using BEDtools.

### Linear mixed model regression analysis

Features per BOCA or standard WRE peaks (as SAF format) were counted using featureCount and normalized to CPM (see above). Outputs were processed in Python (v.3.9.21, JupyterLab). Genotype was encoded as a binary variable and time as a continuous covariate. Global trends (thick lines) were estimated with statsmodels (v.0.14.4) using a linear mixed-effects model of the form: signal ∼time +genotype +time:genotype +(1 ∣ peak); where peak identity was included as a random intercept to account for baseline differences between peaks. The reported slope corresponds to the fixed-effect estimate of time (WT) or the sum of the time and interaction terms (KO).

### Data visualization

Data processing for visualization and plotting was performed in Python (v.3.9.21) within a JupyterLab environment using the *pandas (v*.*2*.*2*.*3), NumPy (v*.*2*.*0*.*2), seaborn (v*.*0*.*13*.*3)*, and *matplotlib (v*.*3*.*9*.*4)* libraries. Signal tracks were visualized using IGV (v.2.19.1 ^58^).

