## Supplementary Figures for "Enhancer pioneering activity of Wnt/β-catenin signaling"

### Supplementary Information

#### Supplementary Figure 1

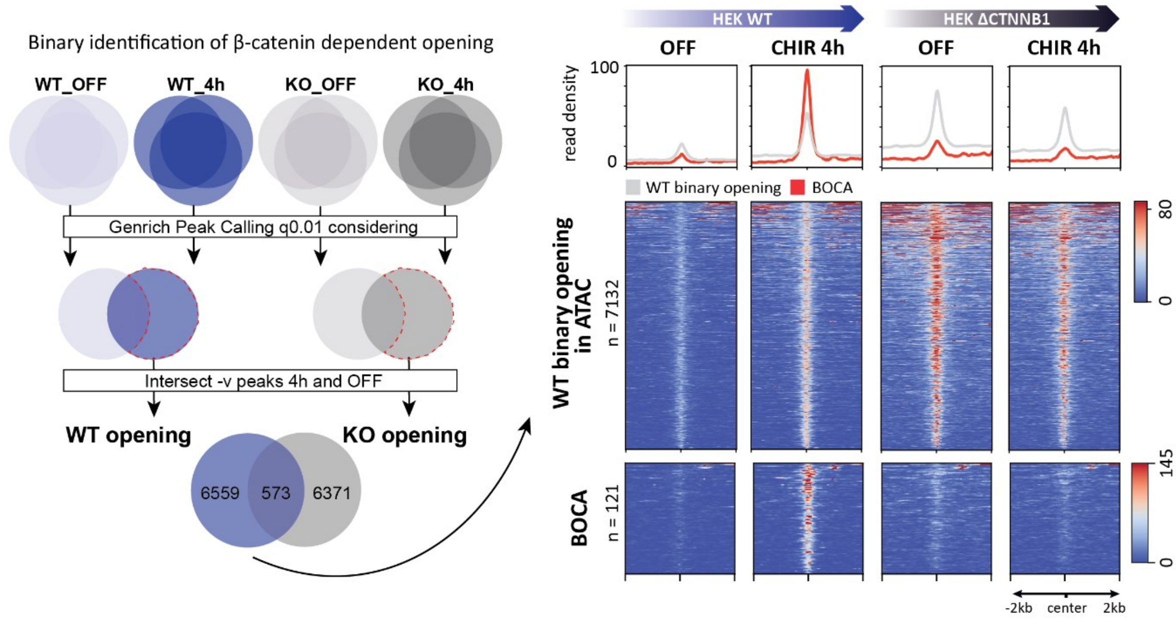


**Right**: Conventional binary identification of opening events: The replicate concordance from the OFF conditions overlapped with the replicate concordance of the CHIR 4h conditions, which leads to 7132 sites that are uniquely open in HEK WT upon CHIR (red dotted line). However, only ~92% of these opening events overlap with the corresponding KO (*ΔCTNNB1*) opening events. When visualizing the ATAC-seq signal across sites (**Left**), the majority of regions that increase in accessibility in WT cells upon CHIR are already accessible in the HEK *ΔCTNNB1* cells. Thus, we decided to follow a more quantitative, stringent approach, that truly captures all β-catenin dependent opening events with closed chromatin in WT OFF, *ΔCTNNB1* OFF and *ΔCTNNB1* 4h, and significant increase of ATAC-seq signal in WT 4h only, as described in Figure 1. If the global shift towards baseline accessible chromatin under Wnt-OFF conditions in *ΔCTNNB*1 cells reflects a clonal artifact or represents a biologically meaningful consequence of the *CTNNB*1 loss remains to be determined.

#### Supplementary Figure 2

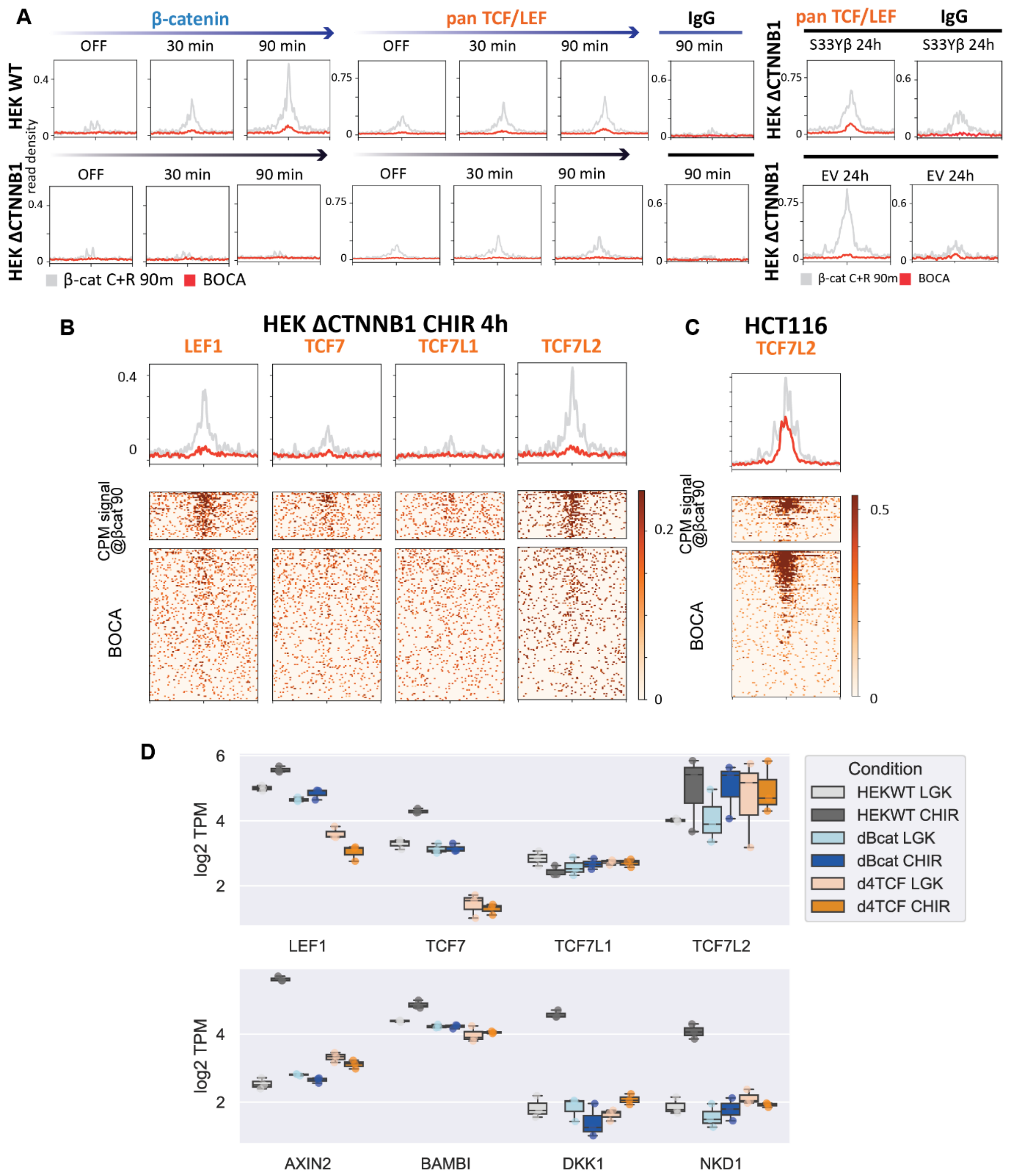


**A.** Average plots comparing CPM normalized CUT&RUN signal of β-catenin, panTCF/LEF and an IgG control, relative to conventional WREs (gray) and BOCA sites (red) ±2 kb around peak centers. Signal corresponds to samples shown in Figure 2A and 2D. **B.** Average plots (top) and heatmaps (bottom) of CPM-normalized CUT&RUN signal generated using antibodies against LEF1, TCF7, TCF7L1, and TCF7L2 individually in HEK *ΔCTNNB1* at 4h CHIR. Although each antibody has been extensively validated for CUT&RUN before, we further confirm that TCF/LEF binding is absent at BOCA sites for all individual factors in HEK *ΔCTNNB1*. This supports the validity of the panTCF/LEF approach used in the main figures where all antibodies were combined. **C.** TCF7L2 shows strong enrichment to BOCA sites in HCT116 cells, which harbor a constitutive Wnt pathway activation, highlighting the functional relevance of these enhancers. **D.** Top: Expression levels (log2TPM) of TCF/LEF family members, showing that LEF1 and TCF7L2 are the most highly expressed factors. This is consistent with our previous finding that these factors exhibit the strongest binding at cWRE in HEK cells, further supporting antibody specificity. Bottom: Expression of four genes (*AXIN2*, *BAMBI*, *DKK1*, *NKD1*) associated with BOCA sites (GREAT). All genes are responsive to Wnt pathway activation, and their expression is reduced upon loss of either panTCF/LEF (orange) or β-catenin (blue). Data from (Doumpas *et al.*, 2019).

#### Supplementary Figure 3

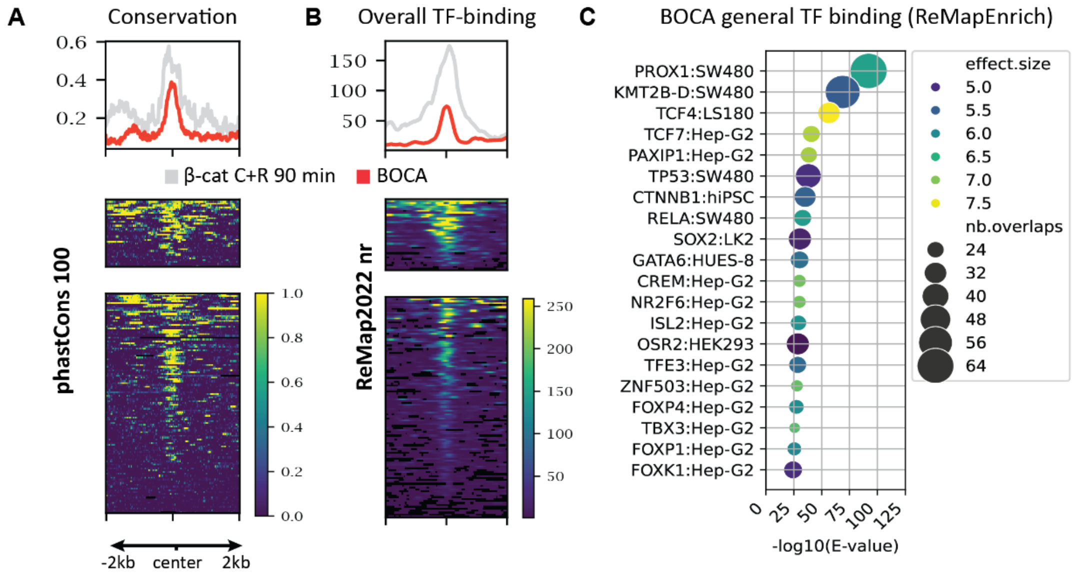


**A.** Sequence conservation across 100 vertebrates at standard WRE and BOCA sites, downloaded from UCSC. **B.** Overall ReMap TF-binding intensities across factors and tissues at standard WRE and BOCA sites. Data downloaded from Remap2022 non-redundant (nr) (Hammal, Langen and Lopez, 2022). **C.** Dotplot showing top 20 of factor:condition combinations that consistently show signal at BOCA sites using the R-package ReMapEnrich (https://github.com/remap-cisreg/ReMapEnrich) (Ch *et al.*, 2020). ReMap complements sequence-based TFs binding predictions by integrating experimentally derived data, systematically mapping TF–DNA associations across the genome using curated datasets.

#### Supplementary Figure 4

| | | Term | Coefficient | Std. Error | p-value |
|---|---|---|---|---|---|
| CBP | βcat 90 | Intercept | 0.159 | 0.0233 | <0.001 |
| | | time | −0.00030 | 0.0002 | 0.138 |
| | | KO | −0.0735 | 0.0158 | <0.001 |
| | | time × KO | 0.00076 | 0.00029 | 0.009 |
| | BOCA | Intercept | 0.1876 | 0.0151 | <0.001 |
| | | time | 0.00105 | 0.00016 | <0.001 |
| | | KO | −0.0514 | 0.0124 | <0.001 |
| | | time × KO | −0.00082 | 0.00023 | <0.001 |

βcat 90: signal = 0.159 + 0.0003(time) − 0.075(KO) − 0.001(time × KO) + (1 | peak)
BOCA: signal = 0.188 + 0.001(time) − 0.051(KO) − 0.00082(time × KO) + (1 | peak)

| | | Term | Coefficient | Std. Error | p-value |
|---|---|---|---|---|---|
| HDAC1 | βcat 90 | Intercept | 0.2238 | 0.0301 | <0.001 |
| | | time | 0.00052 | 0.00026 | 0.051 |
| | | KO | −0.0797 | 0.0205 | <0.001 |
| | | time × KO | −0.00080 | 0.00037 | 0.032 |
| | BOCA | Intercept | 0.2483 | 0.0261 | <0.001 |
| | | time | 0.00267 | 0.00029 | <0.001 |
| | | KO | −0.0862 | 0.0222 | <0.001 |
| | | time × KO | −0.00262 | 0.00041 | <0.001 |

βcat 90: signal = 0.224 + 0.001(time) − 0.08(KO) − 0.001(time × KO) + (1 | peak)
BOCA: signal = 0.248 + 0.003(time) − 0.086(KO) − 0.003(time × KO) + (1 | peak)

**Left:** Table shows individual results from modelling CBP/HDAC1 chromatin association upon stimulation. **Right:** Full function describing the trend line depicted in the main figure for conventional WRE (β-catenin CUT&RUN 90 min CHIR) and BOCA sites, and for CBP and HDAC1.
